# Can SMILES be fragmented into a concatenable ordered sequence of retrosynthetically interesting string block?

**DOI:** 10.64898/2026.08.25.747180

**Authors:** Etienne Reboul, Harish Prabakaran, Jérôme Waldispühl, Antoine Taly

## Abstract

Molecules generated by deep learning models are often difficult to synthesize. Their synthetic accessibility can be improved with automated retrosynthetic analysis, which allows for identifying synthons. However, synthons in a SMILES can be scattered throughout the string depending on the path taken through the molecular graph used to generate the SMILES. We tested whether the ensemble of possible SMILES for a molecule can be used to generate a concatenable ordered sequence of string fragments (blocks) from SMILES that match potential synthons obtained through automated retrosynthetic analysis. We found that exhaustively sampling the SMILES space of a molecule improves the coverage of retrosynthetic breaks. We achieved full coverage of retrosynthetic bonds in string form for 85% of the 1.9 million molecules in the MOSES dataset. Doing so allowed us to test our block SMILES in an unconditional de novo drug design test case with MolGPT and Monte Carlo Tree Search (MCTS). We found that using blocks as an LLM’s token did degrade MolGPT performance due to the curse of dimensionality. However, using the SMILES selected by our blocking algorithm with the default SMILES tokenizer improved the reproduction of physico-chemical properties of samples and also improved uniqueness, novelty, and validity. The MCTS outperforms our MolGPT models in terms of validity and novelty. However, samples generated by the MCTS had physico-chemical properties that were further away from the MOSES baseline than the samples produced by molGPT, with an improved distribution of quantitative estimation of drug-likeness (QED).

## Introduction

De novo Drug Design is the process of creating novel compounds with desired biological and physicochemical property profiles from scratch. This can be achieved using generative chemistry, which consists of using machine learning and deep learning algorithms to generate novel compounds with a predefined set of parameters. Currently, a popular deep learning approach is to reuse model architecture developed by the Natural Language Processing (NLP) community for Large Language Models (LLM). This approach yielded the so-called ‘Chemical Language Models’ (CLM) ^1^ that use molecular string representations such as the Simplified Molecular Input Line Entry System (SMILES) ^2^ or Self-Referencing Embedded Strings (SELFIES). ^3^ A recurring challenge in generative chemistry is both preserving the relevant scaffolds, which are often responsible for the molecule’s pharmacological activity, and ensuring that the molecules are synthetically accessible. One of the solutions to address the latter problem is retrosynthetic analysis. Retrosynthetic analysis is used by organic chemists to plan their chemical syntheses. It consists of the iterative breakdown of a target molecule into synthons. Synthons are hypothetical chemical substructures that, when combined, form the target compound and can also be matched to commercially available reactants. ^4,5^ Chemoinformatics algorithms such as RECAP, ^6^ BRICS, ^7^ and its updated version, R-BRICS, ^8^ aim to automate retrosynthetic analysis. Subsequently, it is possible to represent the scaffold of the molecule as one or a combination of synthons.

Making synthons evident in SMILES is challenging due to how SMILES are generated. SMILES are generated by performing a depth-first traversal of a molecular graph. Thus, the SMILES string generated depends on the starting point and the path taken during the depth-first traversal. SMILES can be understood as a linear, one-dimensional graph representation. This atom-level linearization can distribute information about synthons across distant parts of the string, making it difficult to represent chemically meaningful string fragments. By design, standard SMILES are not the most well-suited to intelligently represent synthons or other chemically relevant blocks in a string.

### SMILES fragmentation

To address this issue, multiple Fragment-based representations derived from SMILES were developed in the past few years. Some examples of such representations include t-SMILES, ^9^ fragSMILES, ^10^ SAFE, ^11^ and SMILES Pair Encoding (SPE). ^12^ The key concept behind these techniques is to decompose the molecular graph into synthons or relevant chemical substructures and assemble the SMILES of each subgraph to form a string. t-SMILES assembles subgraphs as a binary tree, with each ‘branch’ representing a subgraph of the molecule. The SMILES of each subgraph are assembled into a string using the token “&” as a delimiter; the anchor points for each subgraph are represented in SMILES by dummy atoms “*”. The binary tree branching is represented by the tokens “^”. SAFE uses another approach to link SMILES fragments that is based on the permissive grammar of SMILES. Digits in SMILES strings traditionally represent ring closures. However, they can also be used to connect any non-adjacent atoms in a SMILES string that do not form a ring. This approach allows for the representation of a molecule as an unordered sequence of synthons that can be parsed as SMILES by RDKit without any additional post-processing. fragSMILES focuses on two aspects: first, figuring out a way to prevent the multiplication of SMILES of the same fragment, and secondly, the loss of relevant stereochemistry. The first objective is achieved by representing each fragment using its canonical SMILES.

A unique SMILES can be consistently found using a canonicalization algorithm. ^13^ Therefore, the same fragment appearing in different molecules can always be represented by the same SMILES. To take into account various possible connections to a fragment, i.e., chemical bonds to other fragments,anchor points are described by the token *n*, which represents the index of the atom to be used as an anchor. The next point is to explicitly store the absolute conformation of the asymmetric carbon in fragments using stereochemistry symbols: *R* and *S*.

One unexplored avenue of the problem is whether, instead of assembling a set of SMILES fragments, a SMILES can be fragmented into a set of string blocks that coincide with synthons. A SMILES string can safely be cut on a bond token, e.g., “-”,“=”,“#”, that is not between two matching parentheses or digits. For each molecule, a plethora of SMILES may exist, as long as the molecular graph is reasonably complex. We can re-frame our objective as finding one or more SMILES that can be safely fragmented at positions that yield synthons or interesting chemical substructures. Finding an algorithmically optimal way to traverse the molecular graph to identify the best match with synthons is a difficult problem. ^16^ Nonetheless, generating different SMILES for a molecule is relatively cost effective, thanks to the RDKit C++ vectorized implementation of randomizing the path taken to traverse the molecular graph. This enables the generation of a plethora of different SMILES. SMILES annotation for string fragmentation is also efficient because string processing is well optimized, and the time complexity of fragmentation is reasonable, as only a single pass is needed to fully annotate a SMILES.

### work outline

The problem of finding a SMILES that can be safely fragmented into string blocks can be framed as a problem of finding an algorithm to sample SMILES that can be decomposed as synthons. The first part of this study will be dedicated to modeling the maximum number of unique SMILES per number of randomized SMILES generated using the RDKit randomization method. This will allow us to determine whether extensively sampling the SMILES of a molecule is possible and how costly it would be. The second step of this study is to create a library of fragmented SMILES. which we will call SMILES Block, by combining automated retrosynthetic analysis with syntactically safe SMILES. The third and final step will be to run test cases of the use of block SMILES.

## Methods

### data

The fragmentation algorithm described in the ‘Syntactically valid SMILES fragmentation’ subsection was applied to the MOSES dataset, ^17^ which is derived from the ZINC15 database. ^18^ MOSES is constructed from the ZINC15 Clean Leads subset and is curated to represent druglike molecules. The filtering criteria included molecular weights between 250 and 350 Da, a maximum of seven rotatable bonds, exclusion of charged species (e.g., nitrates), restriction to the elements C, N, O, S, F, Cl, Br, and H, and elimination of ring systems larger than eight atoms. The final dataset is divided into the original publications available sets: a training set of approximately 1.7 million compounds, a standard test set of 176,000 compounds, and a scaffold test set of 176,000 compounds, where the scaffolds are distinct from those found in the training set.

### Retrosynthetic analysis with R-BRICS

The automatic retrosynthetic analysis was performed using the R-BRICS algorithm. Briefly, BRICS and revised BRICS use SMILES arbitrary target specification (SMARTS) patterns ^19^ to identify what are referred to as “fragment prototype” and “linkage prototype”. ^7^ Fragment prototypes are defined as the parts that can be broken out of the molecules, while the linkage prototype defines which bonds can be broken to fragment the molecule. The original BRICS algorithm had 16 fragment prototypes and 46 prototype linkages. The updated version of r-BRICS ^8^ includes ten new fragment prototypes and forty-three prototype linkages. Note that for our particular use case, we do not consider prototype linkages and fragments for ring decomposition.

The justification for such a choice is that a ring in a SMILES cannot be broken into one or more parts without causing a syntax error.

### Randomized SMILES generation & modeling

In this subsection, we detail the protocol used to model the number of unique SMILES generated per the number of randomized SMILES. The function used to generate the randomized SMILES is RDKit’s *rdkit.Chem.rdmolfiles.MolToRandomSmilesVect*; as the name suggests, it is a vectorized implementation of the option random in the classic MolToSmiles function from the Chem module.

A simple incremental generation protocol with an early stopping criterion to limit the waste of computational resources is described in algorithm 1 as follows:

**Algorithm 1.**
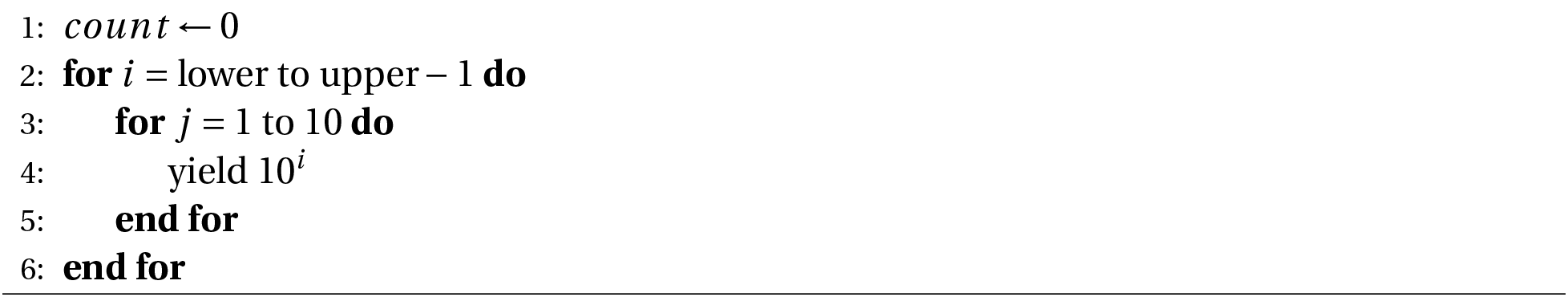
To determine number of randomized SMILES to be generated per iteration.

The lower and upper boundaries for algorithm 1 are 0 and 5, respectively. If the cumulative number of unique SMILES does not increase for 5 consecutive iterations, the generation is stopped early. Thus, the number of randomized smiles generated is a minimum of 5 due to the early stopping criterion, and the maximum is 1 million due to computational and time constraints.

Generating randomized SMILES is a stochastic process that is prone to variability. To take this variability into account, we run the generation protocol in triplicate. The set of unique SMILES from each replica is aggregated into one to maximize the likelihood of capturing all possible SMILES per molecule.

The curve modeling was performed using the python library scipy ^20^ function *curve_fit*. We modeled the number of unique SMILES per number of generated randomized SMILES. We tested all the function listed in table 1.

**Table 1:** Model Functional Forms.

| Model name | Formula |
| --- | --- |
| Square Root | $y = \alpha \sqrt{\beta x}$ |
| Logarithmic | $y = \alpha \log(\beta x)$ |
| Inverse | $y = \frac{\alpha}{\beta + \frac{1}{x}}$ |
| Inverse 2 | $y = \frac{\alpha}{1 + \frac{\beta}{x}}$ |
| Exponential | $y = \alpha (1 - e^{-\beta x})$ |

### Syntactically valid SMILES fragmentation

**Algorithm 2.**
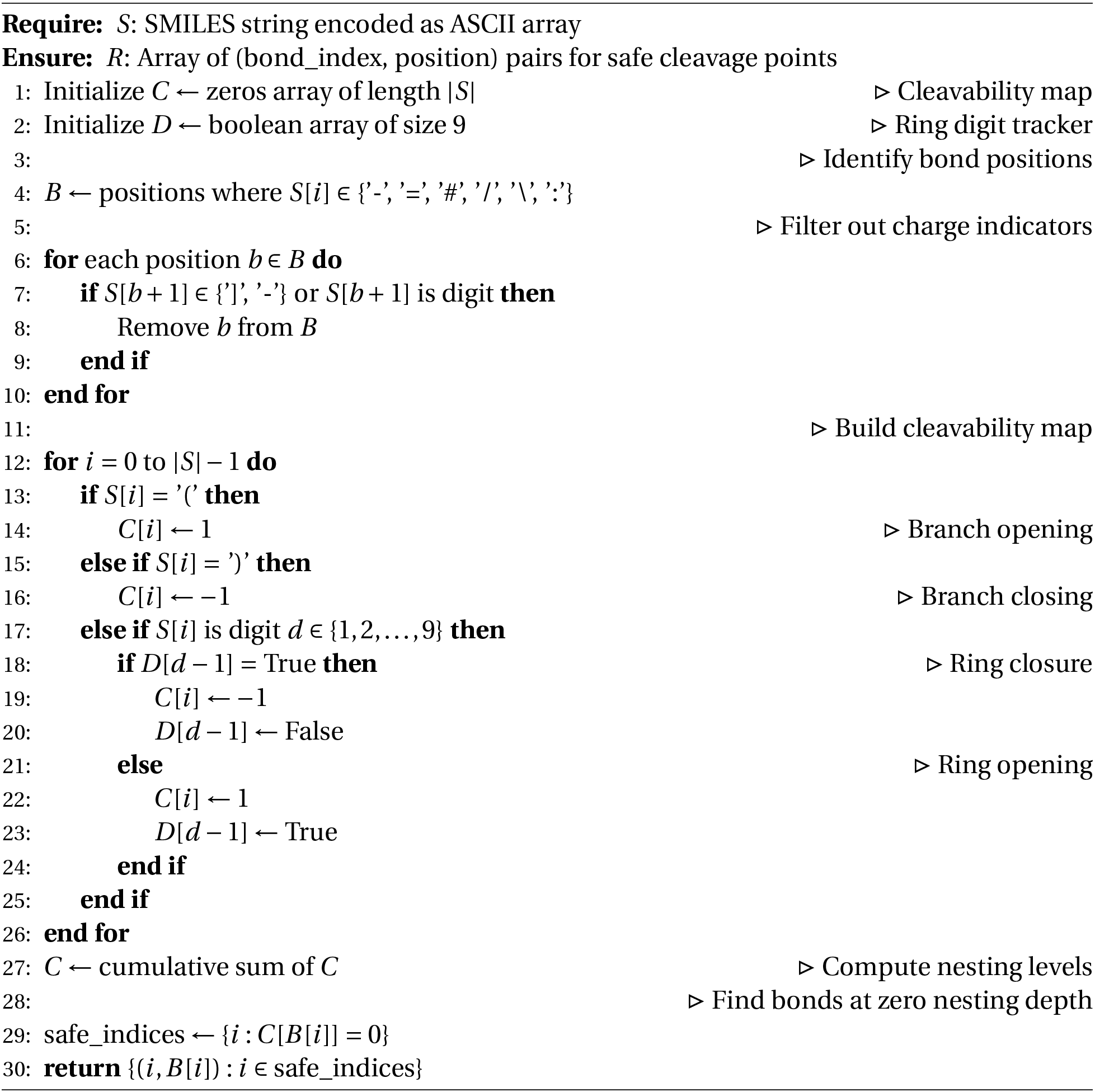
Find Safe SMILES Fragmentation Points.

Our new algorithm identifies safe fragmentation points in SMILES strings by locating chemical bonds that can be cleaved without violating SMILES grammar. The fragmentation algorithm operates by first identifying all bond characters in the SMILES string, then constructing a cleavability map that tracks the nesting level of branches and ring closures. Bonds are considered safely cleavable only when they occur at zero nesting depth, ensuring that fragmentation at these positions preserves the structural integrity of both resulting molecular fragments. The precise SMILES string fragmentation algorithm is described in 2.

### SMILES Blocks generation

Initially, a correspondence is established between retrosynthetic analysis, which identifies bonds that can be retrosynthetically broken, and the SMILES fragmentation process, which detects bonds within a string that, when split, generates two valid SMILES strings. This step is crucial because each Randomized SMILES has its unique atomic and bond indexing, which might differ from the canonical indices used in retrosynthetic analysis. Every bond index can be represented by two atomic indices, indicating the atoms that form the bond. By applying RDKit’s canonical atom index ranking, we can establish a mapping between the current atomic indices and the canonical ones, thereby enabling us to match the bonds identified in both retrosynthetic analysis and string fragmentation.

Following fragmentation, each molecular block is subjected to quality assessment using multiple criteria. Quality metrics included molecular weight, the number of hydrogen bond donors, acceptors, and rotatable bonds, lipophilicity (Crippen logP), and topological polar surface area (TPSA). They are based on an extended version of the rules of 3, to assess each block in each solution:

- Molecular Weight < 300 Daltons
- Number of Hydrogen Donors less than or equal to 3
- Number of Hydrogen Donors less than or equal to 3
- Number of Rotatable bonds less than or equal to 3
- Crippen LogP less than or equal to 3
- Topological Polar Surface Area is less than or equal to 60

We used a simplified version of the fragSMILES blocking system ^10^ to create a unique ID for our blocks that prevents the duplication of string blocks representing the same molecular fragment with the same connections in the same order.

### MolGPT

MolGPT ^21^ is a Transformer-based generative model tailored for de novo molecular design. Built on the Generative Pretrained Transformer (GPT) framework, it employs eight stacked decoder layers, each integrating masked multi-head self-attention and feedforward sublayers with GELU activations. The model is relatively compact, with roughly 6 million trainable parameters, and is optimized using Adam with a learning rate of 6 × 10^−4^. Training is performed over 10 epochs with a batch size of 384, while the learning rate follows a cosine decay schedule, reaching a minimum of 6 × 10^−5^. Cross-entropy serves as the training loss.

During inference, molecules are generated in an autoregressive manner with a sampling temperature of 1.0, enabling a balance between output quality and diversity. Token selection at each step is carried out using a softmax distribution. Within each decoder block, the causal masking of the self-attention mechanism ensures that tokens only attend to previous positions, thereby enforcing the autoregressive property. The attention weights are computed through scaled dot-product attention: ^22^

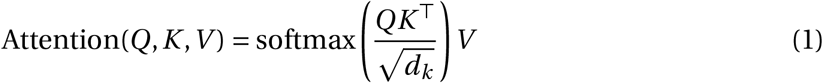

where *Q*, *K*, and *V* correspond to the query, key, and value projections of the input *X*, and *d_k_* represent the dimensionality of the key vectors. This mechanism enables the model to capture long-range contextual dependencies across molecular sequences.

Importantly, the original code of MolGPT uses randomization roughly every two epochs of training to increase the size of the drug-like chemical space explored by generated samples. ^23^ Because this was very difficult to apply to block SMILES, this feature was disabled for all MolGPT models.

### MCTS

Monte Carlo Tree Search (MCTS), ^24,25^ which has attracted growing interest for de novo drug design in recent years, ^26–28^ is used to search the possible combinations of blocks from the block library obtained from MOSES and concatenate those blocks together to form a new molecule. MCTS is an algorithm that balances the exploration of unseen block combinations with the exploitation of known high-scoring paths.Each node in the tree represents a partial molecule, an ordered sequence of block unique identifiers, and carries the statistics (*w*, *v*) representing cumulative reward and visit count, respectively The number of blocks for a new molecule is sampled from a gaussian distribution *N* (*µ*, *σ*^2^). The parameters *µ* and *σ* are obtained by a simple gaussian fit of the blocked SMILES of the MOSES dataset. ^17^ block compatibility is enforced at each step through a revised R-BRICS retrosynthetic tag system ^7^ to increase the probability that combinations of blocks found by the MCTS are synthesizable.

#### Tree Policy

Child selection follows the PUCT (Predictor Upper Confidence Bound for Trees) criterion: ^29,30^

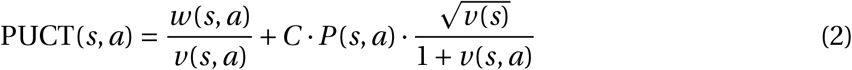

where *s* denotes the current partial molecule, *a* denotes a candidate block extension, *C* is an exploration constant, and *P* (*s*, *a*) is a prior probability over actions.

The prior is set to the empirical conditional probability directly:

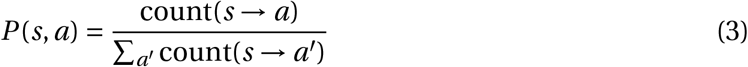

where count(*s* → *a*) is the number of times block *a* was observed as the immediate successor of the last block in the partial sequence *s* across the training corpus.

To prevent the search from collapsing to the prior at the root node, Dirichlet noise ^31^ is injected prior to expansion:

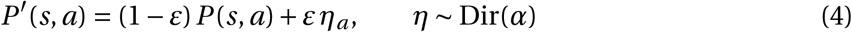

with *ε* = 0.25 and *α* = 0.3, following the AlphaGo Zero parameterization. ^31^

#### Rollout Policy

From each selected leaf node, a complete molecule is assembled by the *default policy*: at each step, the conditional prior (Eq. 3) is used. Sampling continues until a terminal block (end_tag = no_tag) is reached or the target chain length is attained. Compatibility between consecutive blocks is enforced at every rollout step via the revised R-BRICS tag system, so only chemically plausible blocks are considered.

#### Reward Function

A complete molecule assembled during simulation or at a terminal tree node is evaluated by a composite scoring function that combines two soft drug-likeness components:

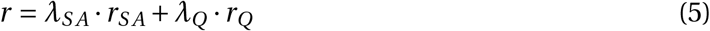

where the two components are defined as follows:

*r_QE_*∈ [0, 1] is a soft synthetic accessibility component ^32^ defined as a linear gradient between the accessibility threshold τ*_Q_*and the minimum possible SA score of 1.0:

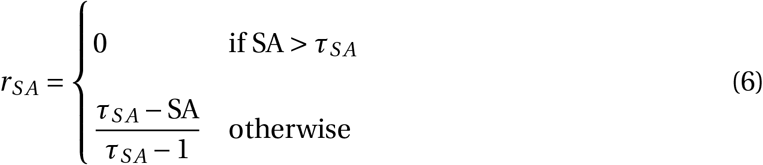

with τ*_Q_* = 4.4 by default.

*r_Q_* ∈ [0, 1] is a soft QED component ^33^ defined as a linear gradient between the drug-likeness threshold *τ_Q_*and the maximum QED of 1.0:

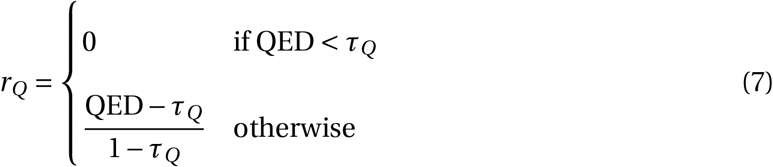

with τ*_Q_* = 0.7 by default, chosen based on the QED distribution of the MOSES training set (Figure 6). Default weights are *λ_S A_* = 0.6, *λ_Q_* = 0.4, and *λ_S A_* +*λ_Q_* = 1, so the maximum possible score is always 1.0.

#### Back-propagation

After a rollout completes and a reward *r* is obtained, the visit count *v* and accumulated reward *w* of every node on the path from the expanded leaf back to the root are updated:

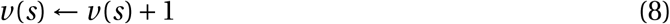

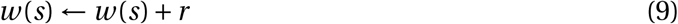

#### Parallelisation and Diversity Control

Root parallelisation ^34^ is used: *N*_workers_ independent MCTS trees are executed concurrently across CPU cores via separate processes. Each process receives a read-only copy of the block library via shared memory (Apache Arrow IPC, zero-copy) and draws its own target chain length independently from the Gaussian prior. All complete molecules scoring above a threshold *τ* are collected from every rollout of every tree.

Runs are organized into sequential waves of *N*_workers_ processes. The search stops at the first wave boundary where

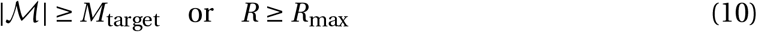

where *M* is the set of unique canonical SMILES collected so far and *R* is the number of completed runs.

Independent trees sharing the same prior distribution tend to converge on the same high-scoring blocks, reducing the diversity of the output. To counter this, a novelty penalty is applied to the block sampling distribution between waves. After each wave completes, the occurrence count *c_a_* of each block *a* is tallied across all collected molecules so far. For the next wave, each worker rebuilds its sampling weights using an adjusted log-weight:

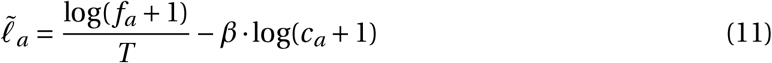

where *f_a_* is the frequency of block *a*, *T* is the temperature, and *β* ≥ 0 is a penalty strength hyperparameter (default *β* = 0.3). The adjusted weights exp(*Ĩ_a_*) are normalized to a probability distribution. blocks that appear frequently in already-collected molecules accumulate higher counts *c_a_* and are therefore downweighted in subsequent waves, steering the search toward less-explored regions of block space. Setting *β* = 0 is equivalent to an unpenalized behavior. The count snapshot used by each wave is fixed at submission time, so all workers within a wave sample from the same adjusted distribution.

## Results

We begin by presenting table 2, which summarizes the different types of SMILES that will be used in this work.

**Table 2:** Overview of the different SMILES used throughout this work.

| Name | Description | Source |
| --- | --- | --- |
| Canonical | A canonical SMILES control variant which is kekulized and with all bonds explicit | MOSES |
| Randomized | SMILES obtained by RDKit’s vectorize randomisation function, that are kekulized and with all bonds explicit | MOSES |
| Well sampled | pool of unique randomized SMILES with different sets of breakable bonds are available, that are tied in maximum compatibility rate with retrosynthetically favorable bonds | randomized SMILES |
| Selected | first entry of the well sampled SMILES list that is selected for fragmentation | well sampled SMILES |
| Block | SMILES fragment obtained by cleaving a selected or canonical SMILES at retrosynthetically favorable bonds | canonical or selected SMILES |

### Generation and modeling of randomized SMILES

One of the critical parts of the pipeline is to ensure that we sample the SMILES space of a molecule sufficiently. Here, we define the SMILES space of a molecule as the enumerated space of all possible SMILES representations for a given molecule. We generated the SMILES data using the protocol described in the subsection Generation and modeling of randomized SMILES from the methods section. Following the aforementioned protocol, we generated approximately 1.25 trillion randomized SMILES, which yielded 281 million datapoints to model the number of unique SMILES per number of randomized SMILES generated for each molecule. To that end, we fitted all the functions described in table 1 for each molecule in MOSES. We found that the highest and best value of *R*^2^ was associated with the Exponential function of formula *y* = *α* 1 − *e*^−^*^βx^* as shown in table 3a. Only one molecule in the entire MOSES dataset had a *R*^2^ inferior to 0.99 with the exponential function. We can note that it only slightly outperforms the two inverse functions that are shown in table 3a.

**Table 3:**
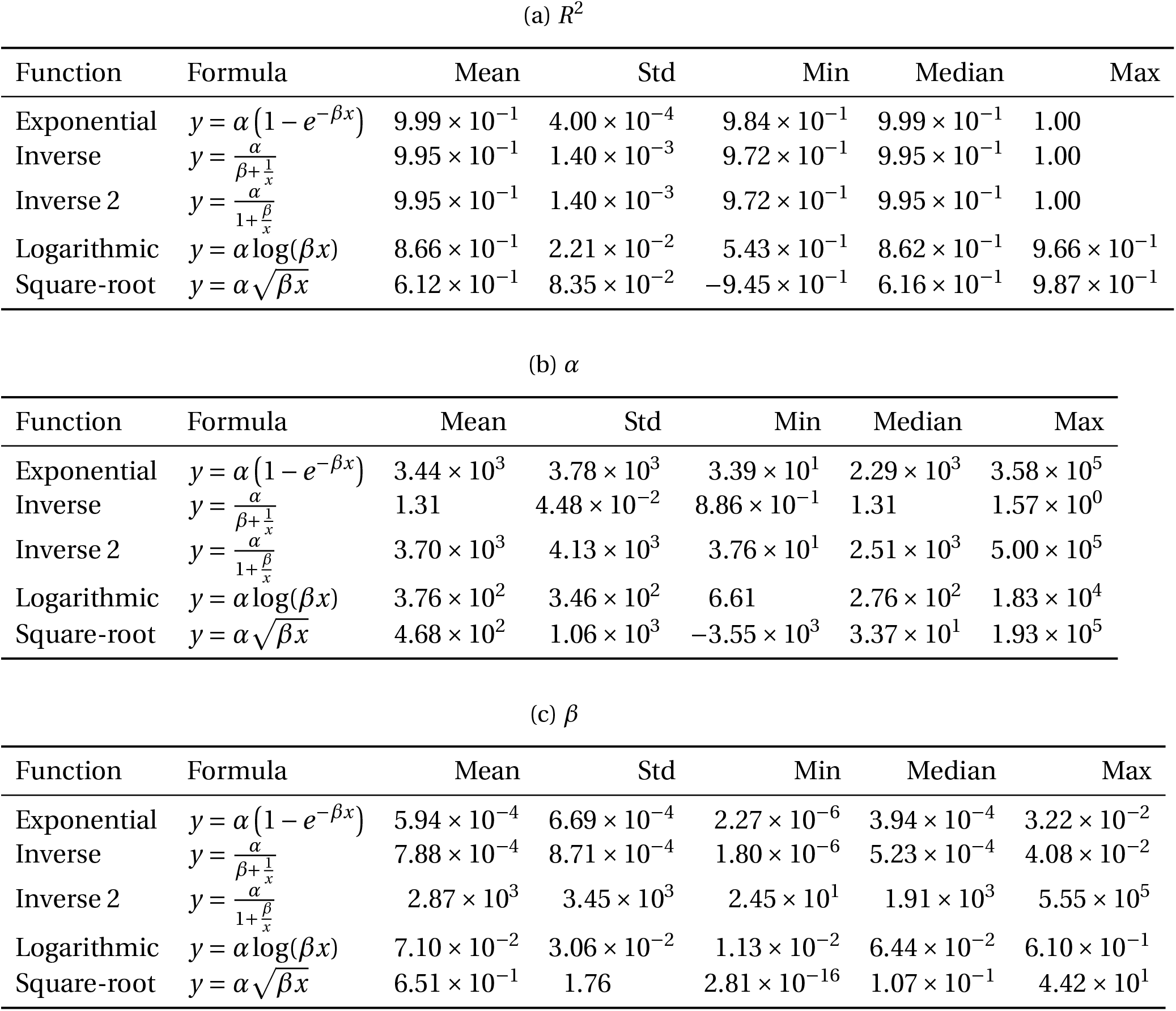
Modeling Function Statistics.

(a) $R^2$
| Function | Formula | Mean | Std | Min | Median | Max |
| --- | --- | --- | --- | --- | --- | --- |
| Exponential | $y = \alpha(1 - e^{-\beta x})$ | $9.99 \times 10^{-1}$ | $4.00 \times 10^{-4}$ | $9.84 \times 10^{-1}$ | $9.99 \times 10^{-1}$ | 1.00 |
| Inverse | $y = \frac{\alpha}{\beta + \frac{1}{x}}$ | $9.95 \times 10^{-1}$ | $1.40 \times 10^{-3}$ | $9.72 \times 10^{-1}$ | $9.95 \times 10^{-1}$ | 1.00 |
| Inverse 2 | $y = \frac{\alpha}{1 + \frac{\beta}{x}}$ | $9.95 \times 10^{-1}$ | $1.40 \times 10^{-3}$ | $9.72 \times 10^{-1}$ | $9.95 \times 10^{-1}$ | 1.00 |
| Logarithmic | $y = \alpha \log(\beta x)$ | $8.66 \times 10^{-1}$ | $2.21 \times 10^{-2}$ | $5.43 \times 10^{-1}$ | $8.62 \times 10^{-1}$ | $9.66 \times 10^{-1}$ |
| Square-root | $y = \alpha \sqrt{\beta x}$ | $6.12 \times 10^{-1}$ | $8.35 \times 10^{-2}$ | $-9.45 \times 10^{-1}$ | $6.16 \times 10^{-1}$ | $9.87 \times 10^{-1}$ |

(b) $\alpha$
| Function | Formula | Mean | Std | Min | Median | Max |
| --- | --- | --- | --- | --- | --- | --- |
| Exponential | $y = \alpha(1 - e^{-\beta x})$ | $3.44 \times 10^3$ | $3.78 \times 10^3$ | $3.39 \times 10^1$ | $2.29 \times 10^3$ | $3.58 \times 10^5$ |
| Inverse | $y = \frac{\alpha}{\beta + \frac{1}{x}}$ | 1.31 | $4.48 \times 10^{-2}$ | $8.86 \times 10^{-1}$ | 1.31 | $1.57 \times 10^0$ |
| Inverse 2 | $y = \frac{\alpha}{1 + \frac{\beta}{x}}$ | $3.70 \times 10^3$ | $4.13 \times 10^3$ | $3.76 \times 10^1$ | $2.51 \times 10^3$ | $5.00 \times 10^5$ |
| Logarithmic | $y = \alpha \log(\beta x)$ | $3.76 \times 10^2$ | $3.46 \times 10^2$ | 6.61 | $2.76 \times 10^2$ | $1.83 \times 10^4$ |
| Square-root | $y = \alpha \sqrt{\beta x}$ | $4.68 \times 10^2$ | $1.06 \times 10^3$ | $-3.55 \times 10^3$ | $3.37 \times 10^1$ | $1.93 \times 10^5$ |

(c) $\beta$
| Function | Formula | Mean | Std | Min | Median | Max |
| --- | --- | --- | --- | --- | --- | --- |
| Exponential | $y = \alpha(1 - e^{-\beta x})$ | $5.94 \times 10^{-4}$ | $6.69 \times 10^{-4}$ | $2.27 \times 10^{-6}$ | $3.94 \times 10^{-4}$ | $3.22 \times 10^{-2}$ |
| Inverse | $y = \frac{\alpha}{\beta + \frac{1}{x}}$ | $7.88 \times 10^{-4}$ | $8.71 \times 10^{-4}$ | $1.80 \times 10^{-6}$ | $5.23 \times 10^{-4}$ | $4.08 \times 10^{-2}$ |
| Inverse 2 | $y = \frac{\alpha}{1 + \frac{\beta}{x}}$ | $2.87 \times 10^3$ | $3.45 \times 10^3$ | $2.45 \times 10^1$ | $1.91 \times 10^3$ | $5.55 \times 10^5$ |
| Logarithmic | $y = \alpha \log(\beta x)$ | $7.10 \times 10^{-2}$ | $3.06 \times 10^{-2}$ | $1.13 \times 10^{-2}$ | $6.44 \times 10^{-2}$ | $6.10 \times 10^{-1}$ |
| Square-root | $y = \alpha \sqrt{\beta x}$ | $6.51 \times 10^{-1}$ | 1.76 | $2.81 \times 10^{-16}$ | $1.07 \times 10^{-1}$ | $4.42 \times 10^1$ |

The *α* parameter in the Exponential function provides an estimate of the maximum number of unique SMILES possible per molecule. This value is very informative, as a large discrepancy between an *α* and the number of unique SMILES found at the end of the generation would indicate that the SMILES space was under-sampled. We computed the ratio between the number of unique SMILES recorded during the generation protocol and the fitted *α* values from the Exponential function. The distribution of those ratios is presented in Figure 2. We found that the agreement between the estimated and experimental value of the number of unique SMILES per molecule is excellent, as 99.33% of the ratios are comprised between 1.00 and 1.02. Only four molecules in the entirety of the MOSES database had a ratio below 0.99, with the lowest recorded ratio being 0.946.

**Figure 1:**
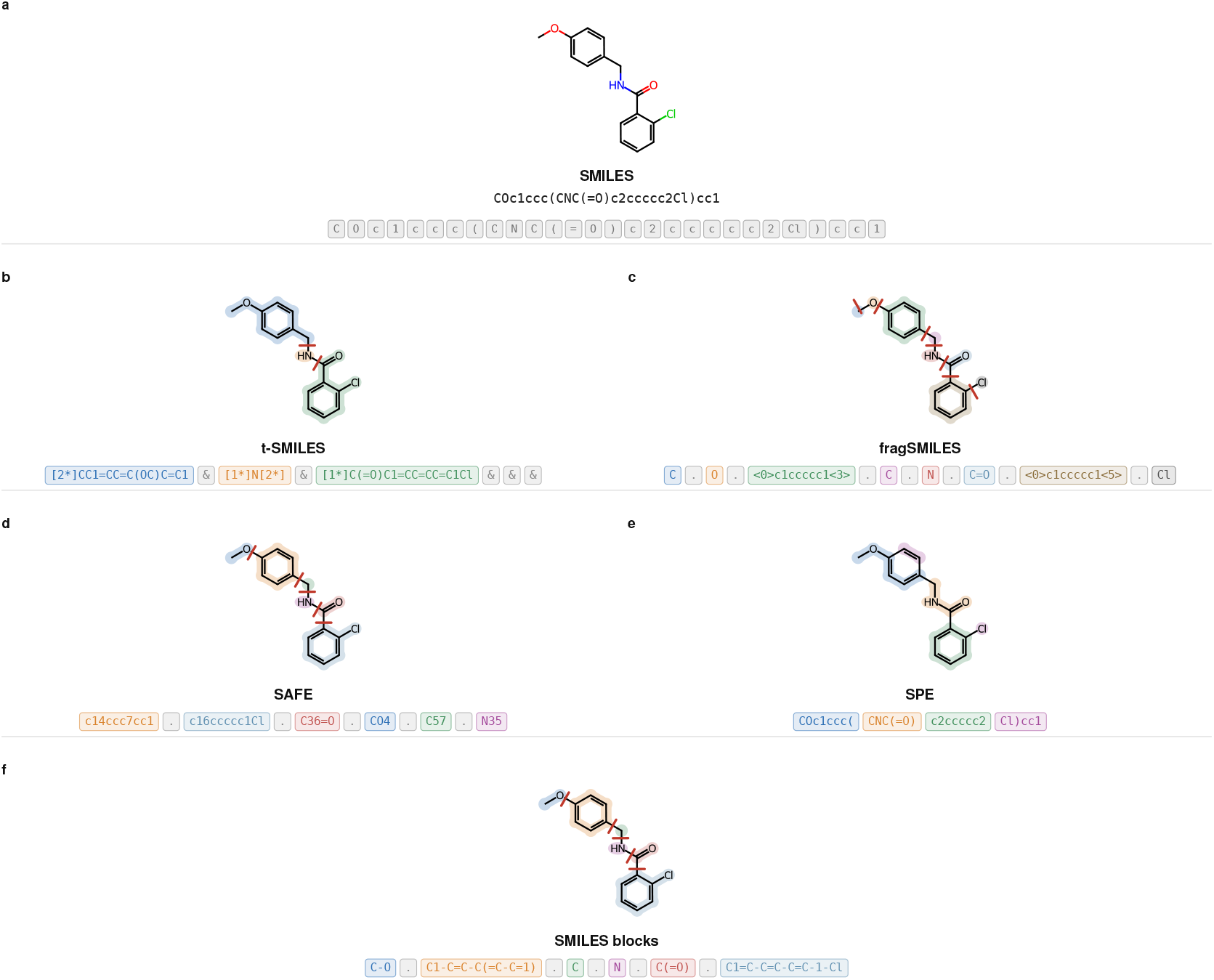
Fragment-based SMILES notations applied to 2-chloro-*N*-(4methoxybenzyl)benzamide.(a) Reference SMILES string and its atom-level tokenisation(b) t-SMILES, ^9^ which fragment molecular graph into a binary tree (c) fragSMILES ^10^ cleaves every exocyclic single bond and writes fragment as canonical SMILES tagged with their attachment points (<i>).(d) SAFE ^11^ fragments the molecule with BRICS and write fragment as independent SMILES joined with ‘.’ and connected via matching ring-closure digits. (e) SMILES Pair Encoding (SPE), ^12^ tokenization base on Byte Pair Encoding (BPE) ^14^ using a database (Chembl ^15^) as reference. (f) SMILES block, our new string based fragmentation methods. In (b)-(d), colour indicates fragment membership and red tick marks show the bonds cleaved to obtain that fragmentation.

**Figure 2:**
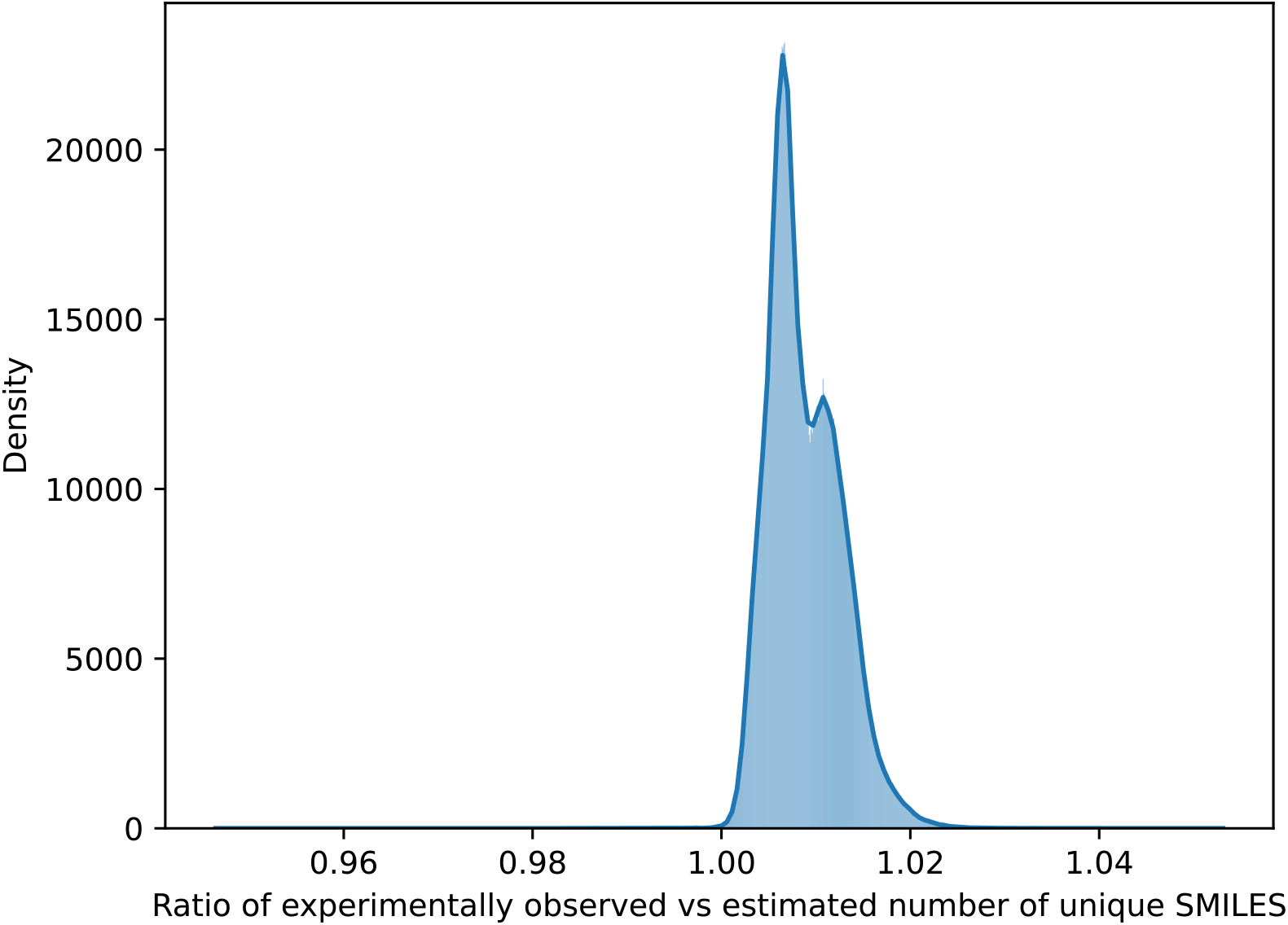
Distribution of the ratio of experimentally observed vs theoretical number of unique SMILES for the exp function

Overall, the modeling of the number of unique SMILES per number of randomized SMILES generated was successful using an Exponential function of the formula: *f* (*x*) = *α*× 1 − exp(−*β* × *x*).

The analysis of the *R*^2^ value and the ratio between the estimated and experimental values of the unique number of SMILES per molecule proved that the fit to the data was of good quality. Thus, we can confirm that we can extensively sample the SMILES space of the molecule to find a SMILES that can potentially be decomposed into synthons. We found that the MOSES dataset of 1.9 million canonical SMILES could be extended when enumerated to roughly 6.72 billion SMILES.

### SMILES fragmentation

Using the results of the modeling experiments, we have obtained an extensive set of unique SMILES for each molecule in MOSES. We sorted the unique set of SMILES using the memory score, a heuristic introduced in our previous paper. ^35^ The memory score is a simple metric that rapidly assesses the complexity of different SMILES for the same molecule. We sorted the SMILES from simplest to most complex. Each SMILES was assessed by our string fragmentation algorithm, and only the simplest SMILES that allowed us to retrieve the maximum retrosynthetically breakable bonds that are also safely breakable in SMILES were kept.

To provide a meaningful reference, we also compute the breakable bond of canonical SMILES derived from MOSES. This will serve as a reference to prove that exhaustive sampling of the SMILES space of a molecule improves the string fragmentation compatibility with the synthon found by R-Brics.

We then proceeded to the retrosynthetic analysis of MOSES and the matchmaking between the breakable components in the SMILES strings and the retro-synthetically breakable bonds. For each molecule, if multiple SMILES strings with different sets of breakable bonds are available, we select the one that has the most complete match to retrieve the most retro-synthetically breakable bonds. Those SMILES will be referred to as well sampled SMILES in the rest of this article. For canonical SMILES, such a selection is not needed because, by definition, there is only one canonical SMILES per molecule.

We first eliminated the molecules that did not yield any results from the retro-synthetic analysis. We found 8077 molecules in MOSES that could not be decomposed by R-BRICS without ring decomposition (cf. subsection Retrosynthetic analysis with R-BRICS), this represents roughly 0.4% of MOSES entries.

We compared the distribution of the retro bond ratio between canonical SMILE and well sampled SMILES, as shown in figure 3. Retro bond ratios are the matched retro-synthetically breakable bonds that are also breakable in a string, divided by the total number of retro-synthetically breakable bonds. Simply put, a bond ratio of 1 means that all retro-synthetically breakable bonds can also be broken in a string without a syntax violation. Conversely, a bond ratio of 0 means that no bond could be matched in the string.

**Figure 3:**
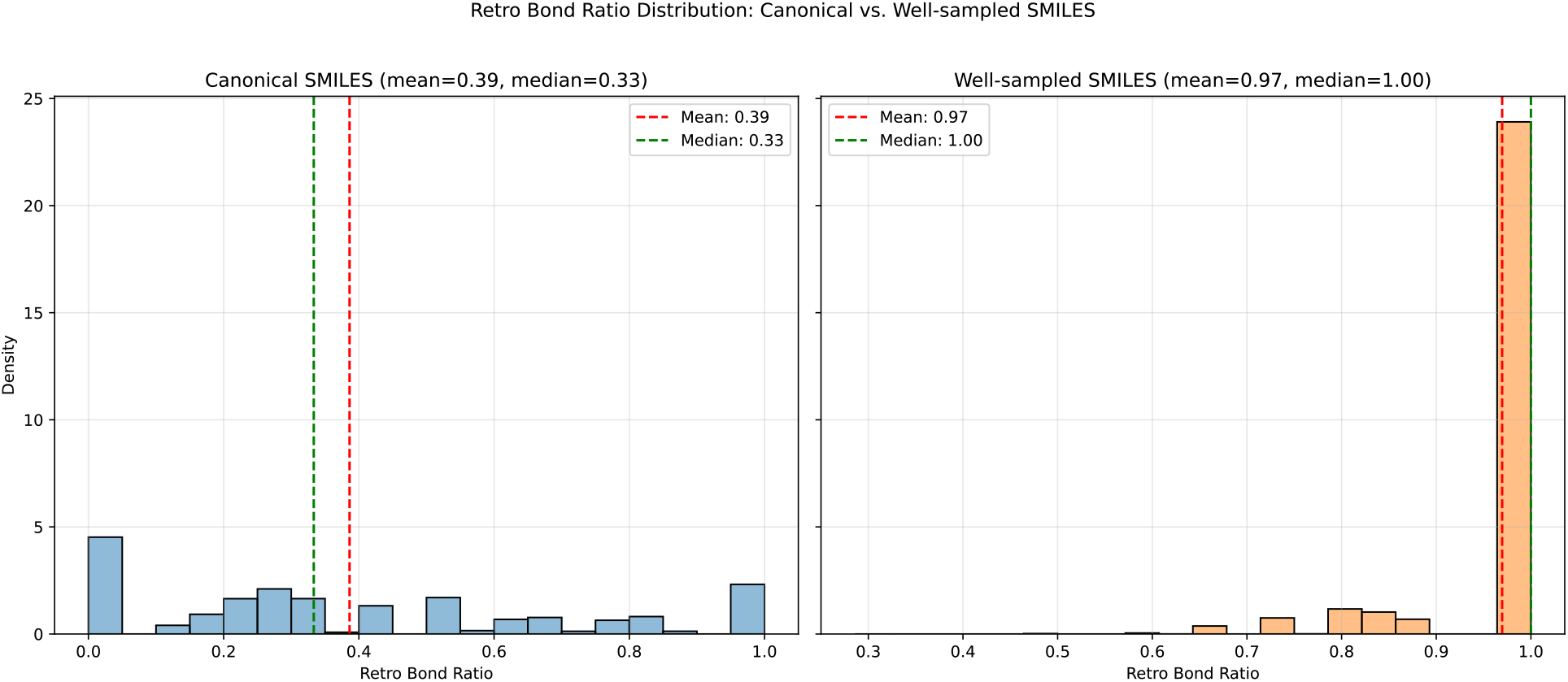
Comparison of retro bond ratio between well sampled SMILES and canonical SMILES. The retro bond ratio represents the success ratio of matches between retrosynthetically breakable bonds and breakable bonds in the string.

There is a drastic difference in the distribution of the bond ratio between canonical SMILES and well sampled SMILES. well sampled SMILES outperform canonical SMILES, with a much higher value of the retro-bond ratio. The most telling number is that around 85% of well sampled SMILES have a retro bond ratio of 1.0, whereas only roughly 15% of canonical smiles have a perfect ratio.

This translates to the fact that an overwhelming majority of well sampled smiles can be fragmented into interesting synthons, whereas only a few canonical smiles can.

We subsequently focused our analysis on the well sampled smile results. We found that most SMILES from MOSES can be divided into 5 to 6 blocks, as shown in Figure 4.

**Figure 4:**
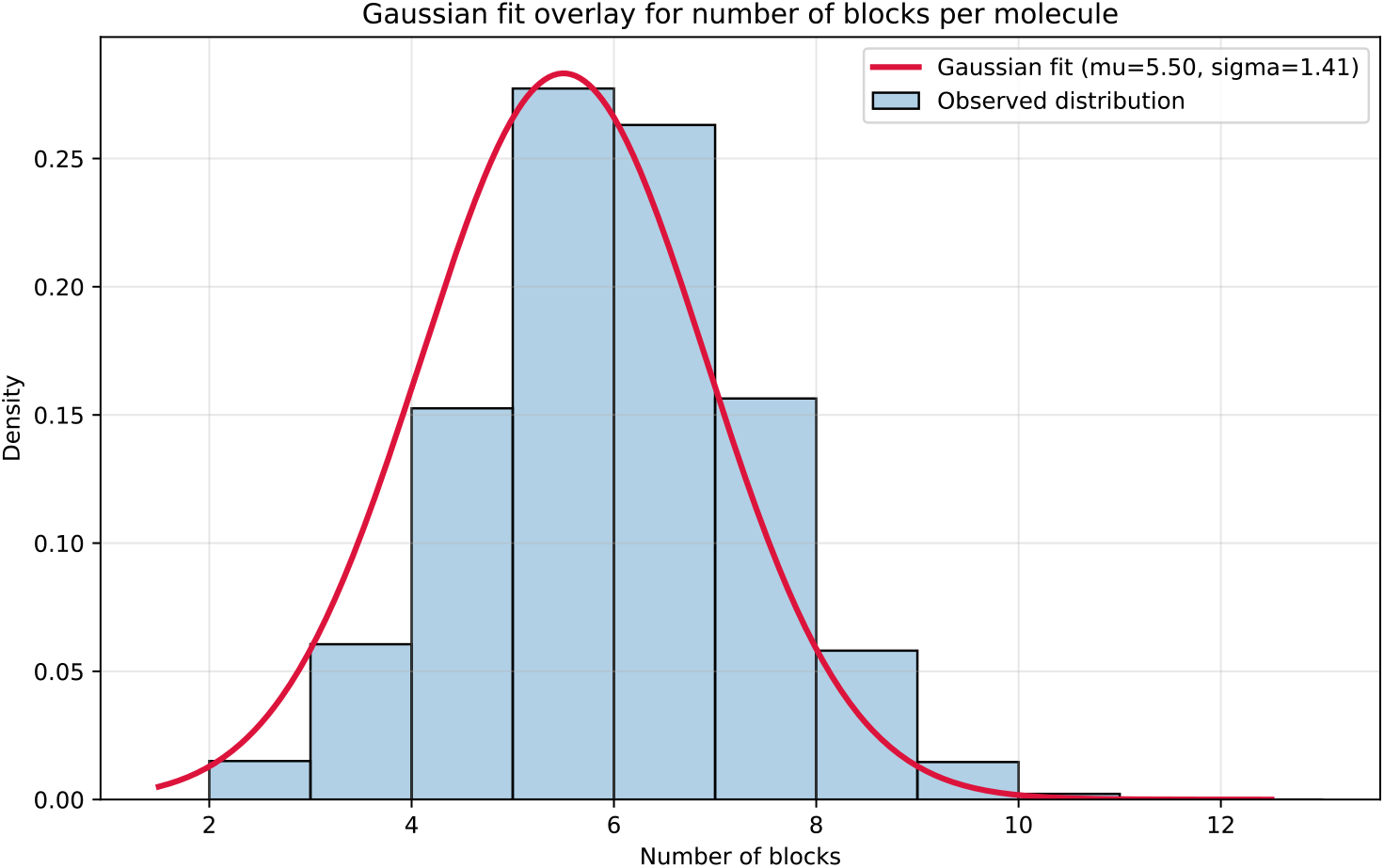
Number of blocks per SMILES for the 1.9 M molecules in MOSES. A Gaussian fit is overlaid as a red curve.

This yielded a library of 119,645 unique blocks derived from the successful fragmentation of well sampled smiles (99.6% of the MOSES dataset). The block distribution is extremely imbalanced, with the top 10 blocks accounting for 46.4% of all block occurrences, and the top 1% of blocks accounting for 88.3% of all block occurrences. The details of the first 12 blocks with the most occurrences in the dataset are shown in Supplementary Information (SI) Figures 1. Additionally, 49.0% of unique blocks occur only once.

We then assessed the quality of those blocks using the extended rules of 3 outlined in the methods section. We found that roughly 73% of the molecules in MOSES could be represented exclusively by blocks that followed every condition listed in the filtration rules. However, only 58.47% of the 119,645 unique blocks pass all the filters from the extended rules of 3.

However, we should note that the rules of 3 and derivatives are more guidelines than imperatives. This means that most of the blocks do not comply with all the outlined rules but are still chemically reasonable. To better understand the quality of the blocks, we decided to plot the distribution of each metric used for filtration across all the blocks, as shown in Figure 5. The two metrics that prevent the most blocks from being considered pure are TPSA and the number of hydrogen donors, with acceptance rates of only 78.7% and 72.4%, respectively.

**Figure 5:**
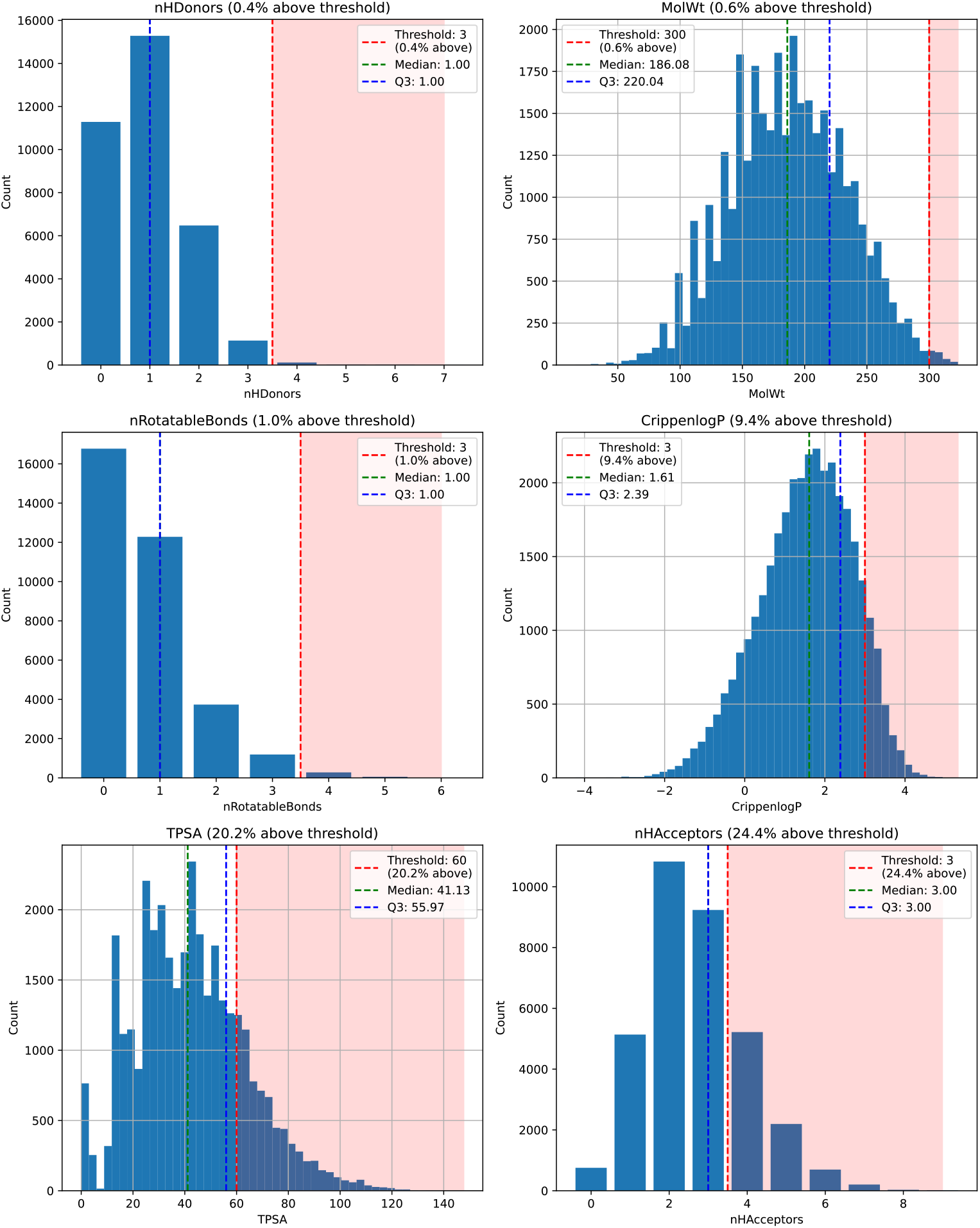
Distribution of the six extended Rule-of-Three properties across the unique well sampled SMILES blocks: molecular weight (MolWt), number of hydrogen bond donors (nHDonors), number of hydrogen bond acceptors (nHAcceptors), number of rotatable bonds (nRotatableBonds), calculated octanol-water partition coefficient (CrippenLogP), and topological polar surface area (TPSA). In each panel, the shaded red region marks values above the extended Rule-of-Three threshold.

Moreover, twelve blocks were randomly selected that did not adhere to one or more of the filtration rules of the extended rule of three. Upon visual inspection, their quality was also deemed reasonable, as shown in SI figure 2.

### Use cases

We then evaluated whether using SMILES blocks could improve performance on de novo drug design tasks, first in an unconditional generation setting with the MolGPT model, ^21^ and subsequently using a Monte Carlo Tree Search (MCTS) algorithm.

#### MolGPT

MolGPT models were trained for 10 epochs on the block SMILES, RDKit canonical SMILES. The training for the MolGPT models on an L40S GPU took approximately 3 and 1 days, respectively. The training took RDKit canonical SMILES; MolGPT uses the standard tokenizer, whereas block MolGPT uses blocks as tokens. To train our model, we used the default settings provided in the source code from the original publication by Bagal *et al*, ^21^ except for the SMILES randomization as detailed in the corresponding methods subsection. From each model obtained after the 10th epoch, we sampled 300k samples. The sampling took 3 hours for the canonical models and a little under 6 hours for the block SMILES MolGPT. We began our analysis by assessing the validity, uniqueness, and novelty of our samples. the results are compiled in table 4.

**Table 4:** Generation metrics for MolGPT variants and block MCTS on 300k samples. Uniqueness is computed over valid molecules; novelty over unique molecules.

| Method | Validity (%) | Uniqueness (%) | Novelty (%) |
| --- | --- | --- | --- |
| MolGPT canonical SMILES | 99.4 | 97.2 | 75.9 |
| MolGPT block SMILES | 91.3 | 82.4 | 88.1 |
| MolGPT selected SMILES | 99.6 | **98.7** | 83.2 |
| block MCTS | **100.0** | 92.5 | **94.9** |

The first verification we conducted was comparing the generation metrics and chemical properties of samples generated by the canonical SMILES MolGPT model from this study with the results from the original publication by Bagal *et al*. ^21^ and a replica model in our previous paper. ^35^ We report that removing the randomization feature did not degrade the generation metrics, as shown in table 4 and the reproduction of chemical properties, as shown in Figure 6. We next analyzed the samples produced by Block SMILES MolGPT and RDKit canonical SMILES MolGPT. Our results indicate that RDKit canonical SMILES MolGPT generally performs better than Block SMILES MolGPT with respect to uniqueness and validity, but not with respect to novelty. A likely reason for the poorer validity and uniqueness of Block SMILES is the curse of dimensionality. The Block SMILES vocabulary comprises approximately 119,645 distinct blocks, whereas the standard RDKit canonical SMILES vocabulary contains only 23 tokens, which amounts to a difference of four orders of magnitude. This very substantial increase in vocabulary size may explain the weaker performance. Furthermore, we demonstrate that the physico-chemical properties of molecules generated by Block SMILES MolGPT deviate more from those in the MOSES dataset than do the properties of molecules generated by RDKit canonical SMILES MolGPT, as illustrated in Figure 6.

**Figure 6:**
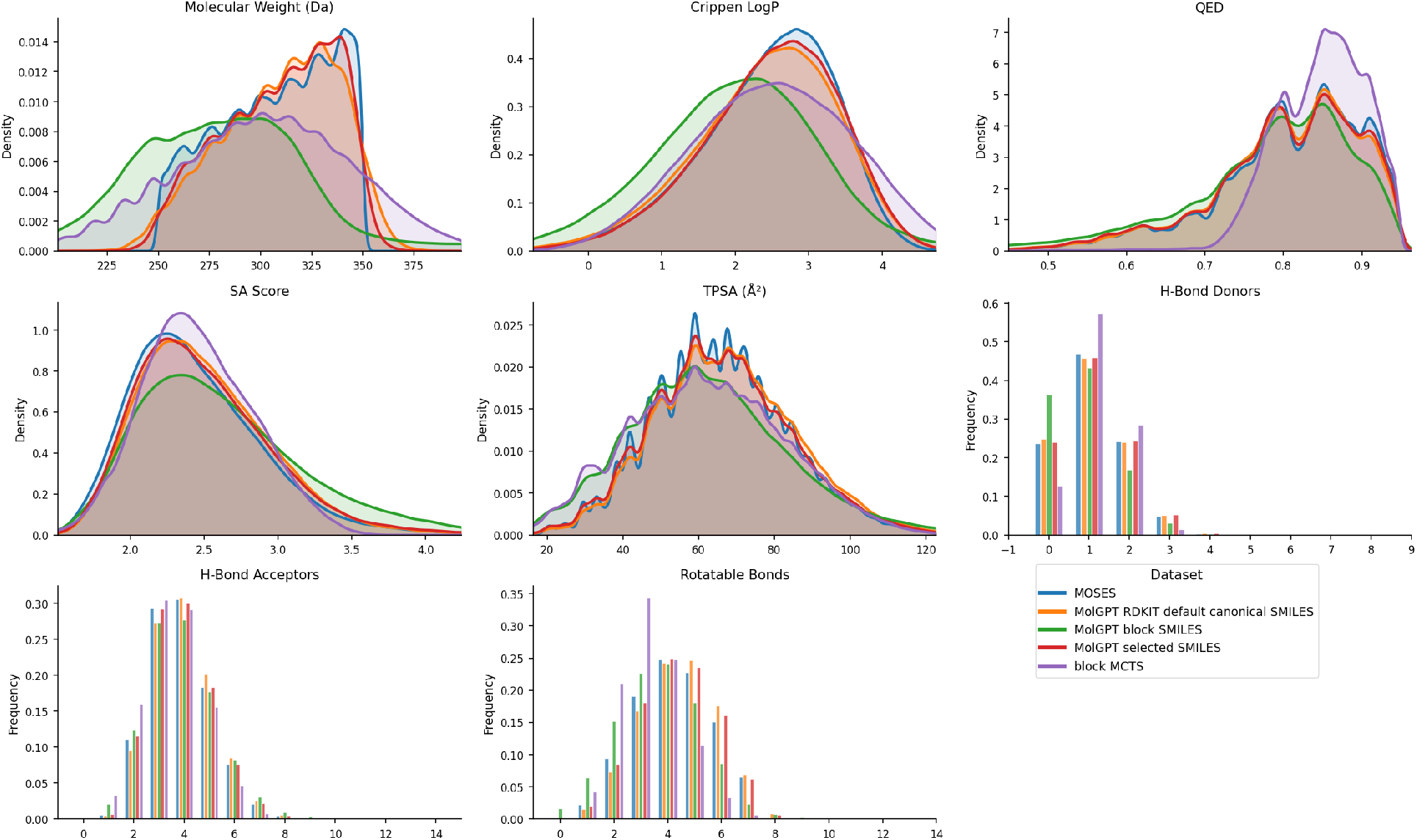
Distributions of molecular properties of 300K samples obtained from MolGPTs canonical SMILES,block SMILES, selected SMILES and the block MCTS overlaid against the MOSES dataset. Properties shown: Molecular Weight, Crippen LogP, QED, SA Score, TPSA, H-Bond Donors, H-Bond Acceptors, and Rotatable Bonds.

Both the SPE ^12^ and SAFE ^11^ approaches employ a Byte Pair Encoding (BPE) tokenization scheme, while t-SMILES ^9^ relies on the standard SMILES tokenizer, extended with additional ASCII characters to represent the new separator. To address our vocabulary size issue, we opted for a solution similar to the latter.

To validate our hypothesis, we trained a MolGPT model on the ‘precursor’ SMILES (selected SMILES) used for block fragmentation, employing the standard SMILES token tokenizer. Compared with the RDKit canonical SMILES MolGPT, the model trained on selected SMILES produced samples with higher validity and uniqueness, as well as a markedly higher novelty rate. In addition, we observed that molecules generated by the selected SMILES MolGPT exhibit physicochemical properties more closely aligned with those in the MOSES datasets.

#### Monte Carlos Tree Search (MCTS)

We generated a little more than 300k samples with the block MCTS which took approximately 10 minutes. MCTS does not require training per se, but the closest preparation akin to training would be the precomputation of connection statistics, which is achieved in under 30 seconds.

Due to the R-BRICS compatibility scheme and the use of connection statistics, the MCTS guaranties 100% validity as non feasible connections between blocks are never considered. Therefore, it outperforms MolGPT models for validity rates, as no reported models have reached 100% validity. It also outperformed MolGPT models in terms of novelty rates. However, it did not outperform MolGPT models for the uniqueness rate, except for the SMILES block model.

Regarding the physico-chemical properties of samples generated with the MCTS, we found that MolGPT models had a distribution closer to the MOSES baseline. Interestingly, the distribution of quantitative estimation of drug likeness (QED) is skewed toward higher values for the MCTS samples, which differs significantly from the original dataset. Another noticeable characteristic of the MCTS samples is that they are more rigid than the other samples, with more compounds having 3 or fewer rotatable bonds.

## Discussion

The principal aim of this study is to determine whether we can generate a concatenable ordered sequence of string fragments (blocks) from SMILES that match potential synthons obtained through automated retro-synthetic analysis.

To that end, we first modeled the number of unique SMILES generated per randomized SMILES for each of the 1.9 million molecules found in the MOSES dataset. We found that the best fit function is an exponential based function described by the formula: *y* = *α* 1 − *e*^−^*^βx^*. The quality of the fit was confirmed by a high *R*^2^ value, with only one molecule not reaching the 0.99 value for MOSES. We were able to confirm the goodness of fit of our exponential models to our experimental data, with the correct prediction of the maximum number of SMILES possible per molecule within a 2% margin of error for more than 99% of the MOSES dataset. This confirmed our ability to sample exhaustively the SMILES space (≈SMILES ensemble) for all molecules in our datasets. The enumeration of possible SMILES for each molecule in the MOSES dataset yields roughly 6.73 billion SMILES. This fully enumerated set of SMILES is going to help future work train Large Language Models (LLMs) on augmented data, and we will likely help improve the quality of generative models. ^23^

We tested which bond could be safely broken in the string for the set of unique SMILES (well sampled SMILES) of each molecule in MOSES. We matched the bonds that can be broken in the string with the breakable bonds determined by retrosynthetic analysis using R-BRICS. We retained only one SMILES per molecule that had the highest match between the breakable bonds in the string and the retrosynthetic analysis. The coverage ratio of retrosynthetic breakage was computed for both the well sampled SMILES and the default RDKit canonical SMILES. We found that exhaustively sampling the SMILES space has drastically improved the coverage of retrosynthetic breakage. Others have very recently proposed TYCHE, a more systematic approach to exploring the SMILES sequence space. ^16^ The improvement brought by TYCHE is mainly on the enumeration of cycles so that blockSMILES would not be modified, given that our algorithm does not allow for breaking cycles.

The suitability of the SMILES blocks for tasks related to Fragment Based Drug Design was assessed using the extended version of the Rule of 3. We found that around three quarters of molecules could be represented with fragments respecting all the filtration criteria. However, roughly 60% of unique blocks passed all the filtration criteria. To further assess the physicochemical properties of our unique blocks, we studied the distribution of all the filtration criteria. We found that although a significant portion of the blocks did not meet all criteria, their physico-chemical properties were still reasonable. This confirms that the rules of 3 and derivatives are more guidelines than absolute rules.

We then tested our block SMILES in an unconditional sampling de novo drug design test case with MolGPT and MCTS. We observed that block SMILES MolGPT underperformed compared to the default RDKit canonical SMILES MolGPT in terms of validity and uniqueness, but not novelty. This result highlights the limitation of the MolGPT model, which is likely explained by the curse of dimensionality due to the four orders of magnitude increase in vocabulary size between RDKit canonical SMILES and Block SMILES. To confirm this hypothesis, we used the ‘precursor’ SMILES without fragmentation, i.e. using the traditional tokenizer. Using those selected SMILES to train a new MolGPT model allowed us to improve the validity, uniqueness, and novelty of the sampled generation. We also analyzed the physico-chemical properties of the samples generated by our different models. We found that the chemical profiles for RDKit canonical SMILES and selected SMILES were closely related to the molecules found in the MOSES training set. As expected, the block SMILES samples generated by MolGPT were further away from the baseline, with selected SMILES samples closer to the original baseline.

We then proceeded to use our MCTS to generate samples using the block SMILES representation. One benefit of these generative methods is that they do not require expensive acceleration hardware, such as high-end GPUs needed for LLMs and neural networks, and can run on a CPU only. In future developments, it may reduce the current reliance on high-performance computing and enable de novo generation on affordable computers or laptops. It is also faster than LLMs by several orders of magnitude, since it does not require any actual training, and the most comparable preparation step can be completed in less than 30 seconds. Inference is also faster and can be run on standard CPUs in tens of minutes, whereas MolGPT models require several hours for 300,000 samples. An advantage of using block SMILES with an MCTS is that it ensures the generation of only syntactically valid SMILES since the R-BRICS scheme prevents block connections that would violate valence rules. However, compared to MolGPT, MCTS was less effective at reproducing the baseline distribution of physico-chemical properties in samples, although its QED distribution was shifted toward higher values. This indicates that the samples generated by MCTS are, on average, more “drug-like” than those produced by MolGPT. This study further confirms the continued interest in the field for MCTS in de novo design, especially in drug design. ^36–40^

## Conclusion

In conclusion, the first scientific contribution is that we demonstrated that an overwhelming majority of samples in the MOSES dataset (80%) can be fragmented into a concatenable ordered sequence of retrosynthetically interesting string blocks. The second scientific contribution of this study is a fully enumerated set of MOSES, freely available on zenodo (cf. Data and software availability section) under the permissive license Creative Commons Attribution 4.0 International (CC BY 4.0). The third scientific contribution is that using block or string representation, which drastically increases the size of the vocabulary, is likely to degrade the performance of MolGPT models. However, the particular ordering of SMILES needed to fragment them into blocks is still valuable to MolGPT models, as it is correlated with an increase in generation metrics and better fidelity to the distribution of physicochemical properties. Lastly, MCTS continued to show promise as a simple alternative to complex neural networks such as MolGPT models.

## Authors Contribution

Etienne Reboul: Software Methodology (Blocks SMILES), Investigation, and Writing Review and Editing Harish Prabakaran: Software (MolGPT code adaptation), Investigation, and Review Antoine Taly: Funding acquisition, Supervision, Review, and Editing.

## Generative AI Usage Statement

Claude Sonnet 4.6 assisted in designing the MCTS module and drafting the methods section, with all code and statements subsequently reviewed by human authors. Overleaf’s Writefull AI was used to rephrase paragraphs.

## Data and Software Availability

The code for smiles blocks is available at: https://github.com/EtienneReboul/smiles_blocks The fork repository of MolGPT from *baghal et al* is available at: https://github.com/EtienneReboul/molgpt the enumerated SMILES of the MOSES dataset is available at https://zenodo.org/records/18874641 the rest of the results data is available at https://zenodo.org/records/21175959

