## supplementary figures 1 and 2 for "Can SMILES be fragmented into a concatenable ordered sequence of retrosynthetically interesting string block?"

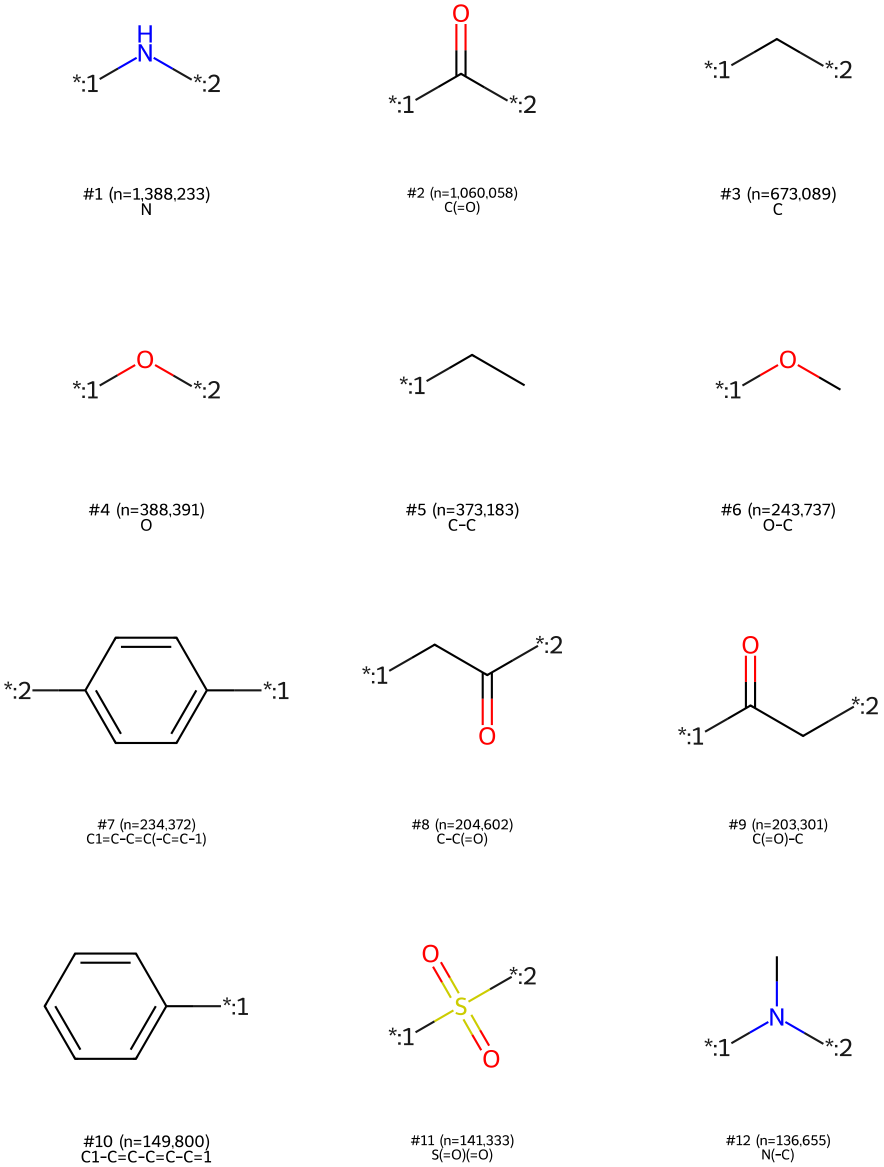


Figure 1: The 12 most represented blocks in the well-sampled MOSES dataset.

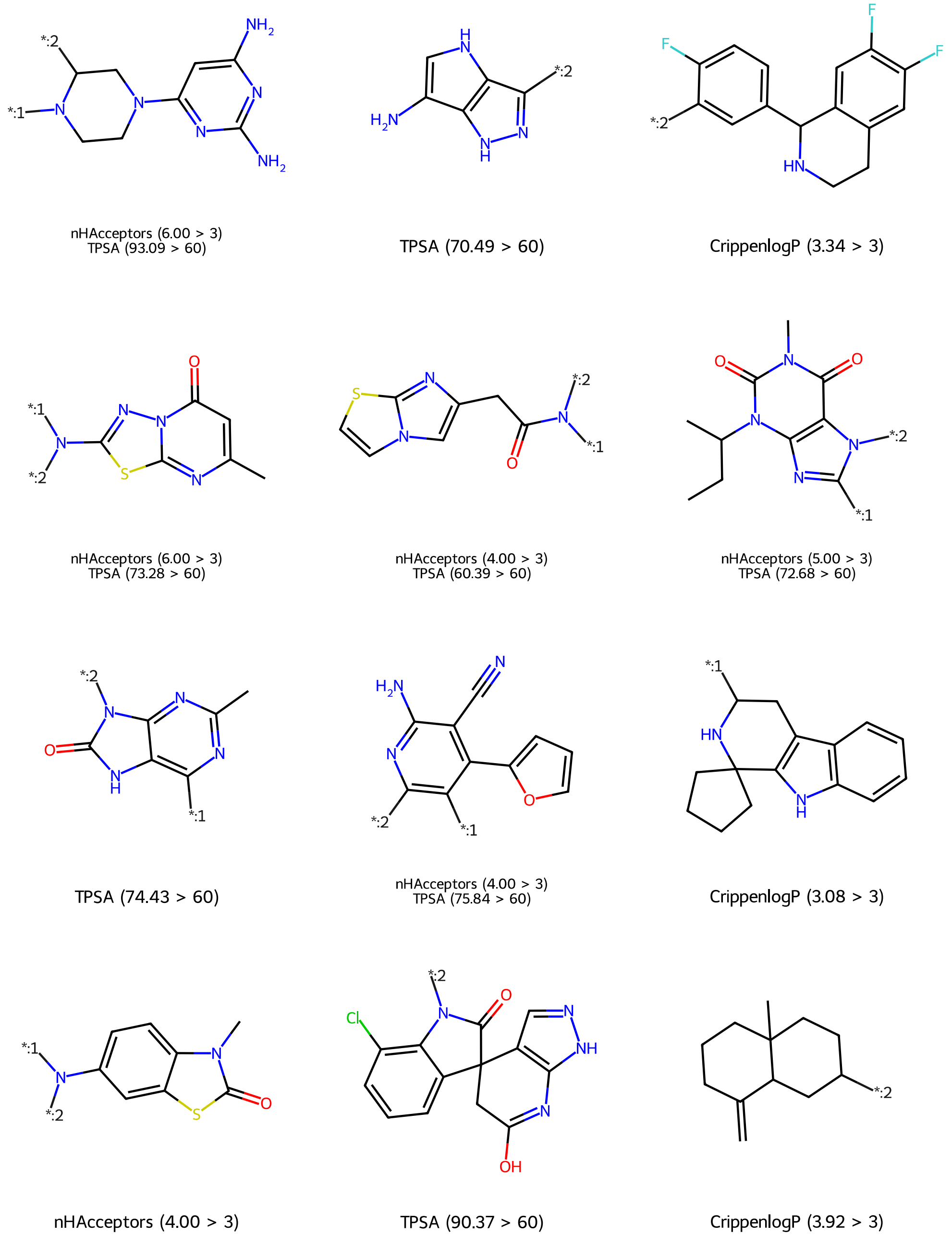


Figure 2: Randomly selected blocks that do no meet one or more criteria from the extended rule of three. The legends of each molecules indicate one or more criteria they did not respect with the threshold value and associated value.
